# Exploring Ventilator-Induced Lung Injury: A Comprehensive Ex-Vivo Study Using Phase-Contrast MicroCT and Atomic Force Microscopy

**DOI:** 10.64898/2026.03.01.708847

**Authors:** Md Motiur Rahman Sagar, Lorenzo D’Amico, Richard T. Deyhle, Ruth Meyer, Luca Fardin, Irma Mahmutovic Persson, Jose Luis Cercos-Pita, Gaetano Perchiazzi, Sarah Köster, Claudia V. Benke, Frauke Alves, Giuliana Tromba, Lars E. Olsson, Sam Bayat, Christian Dullin

## Abstract

Mechanical ventilation (MV) can induce or exacerbate ventilator-induced lung injury (VILI), particularly in mechanically heterogeneous lungs with pre-existing injury. We investigated VILI in a rat model of bleomycin-induced lung injury and compared it with healthy controls using a combined *in-vivo* and *ex-vivo* imaging approach. Previously acquired *in-vivo* data from four-dimensional (4D) phase-contrast synchrotron micro-computed tomography (microCT) and forced oscillation measurements showed increased lung elastance and reduced local acinar strain in bleomycin-induced injured lungs at baseline and after injurious MV. To identify structural and mechanical correlates, we performed automated three-dimensional (3D) pore analysis and atomic force microscopy (AFM) on formalin-fixed, paraffin-embedded lung tissue, complemented by histology and spatial co-registration. *Ex-vivo* analysis revealed pronounced airspace enlargement after injurious MV of healthy lungs, whereas this effect was attenuated in fibrotic lungs. AFM demonstrated region-specific mechanical responses, and correlation analyses linked pore geometry and nanoscale stiffness to *in-vivo* lung mechanics. Spatial analysis further showed colocalization of VILI-associated airspace damage with injured regions. Overall, extracellular matrix remodelling modifies the lung’s mechanical response to injurious MV. This multiscale correlative approach provides mechanistic insight into the interplay between lung injury and VILI and informs ventilation strategies in structurally altered lungs.

## Introduction

Mechanical ventilation (MV) plays a critical role in patient care by supporting gas exchange and alleviating the work of breathing in acute respiratory failure. Unlike natural breathing, which generates negative pressure in the chest upon inspiration, MV generally employs positive pressure. Positive pressure ventilation can induce excessive stress and strain in the lung parenchyma, causing or worsening pre-existing lung injury, a phenomenon known as ventilator-induced lung injury (VILI) [1]. VILI has been extensively studied over the years, leading to refined guidelines for optimal MV settings aiming at limiting tidal volume and plateau pressure [2]. However, the effects of MV on heterogeneous injured lung tissue remain poorly understood, at the microscale. Particularly, the relation between the viscoelastic properties of the respiratory tissues and the local microscopic tissue morphology and mechanical properties remain poorly understood.

In this study, we investigate the effect of injurious MV on microscopic tissue morphology and stiffness in healthy control rat lungs and 7 days post-bleomycin-induced lung injury in rats. The animals were studied *in-vivo* using four-dimensional (4D) phase-contrast micro-computed tomography (microCT) at the European Synchrotron Radiation Facility (ESRF), and quantitative maps of local lung strain were generated [3]. Lung tissue strain analysis using *in-vivo* 4D phase-contrast microCT offers a detailed assessment of lung biomechanics during the respiratory cycle. This approach allows precise measurement of lung deformation at micrometer resolution, providing valuable insights into lung tissue dynamics, stress distribution, and the mechanisms underlying VILI, with potential implications for optimizing MV strategies [4]. Interestingly, we found that bleomycin-induced lung injury, which stiffens the lungs, resulted in reduced local lung tissue strain, both at baseline and following VILI under pressure-controlled ventilation [3]. However, the spatial resolution of *in-vivo* dynamic 4D microCT was too limited to accurately depict lung alveolar morphology.

To further investigate these findings, we present a comprehensive quantification of the local lung structure through automated pore analysis, alongside the correlation with local measurements of lung tissue stiffness using atomic force microscopy (AFM) on formalin-fixed, paraffin-embedded (FFPE) lung specimens from the same cohort of animals.

Three-dimensional (3D) pore analysis is a technique widely used for characterizing porous materials in datasets from 3D imaging techniques, particularly microCT. This method has also been applied in biomedical research, for example, to study changes in trabecular bone [5]. Given the structural resemblance between lung tissue and porous materials, this technique has been successfully used to characterize pathological changes in lung disease models [6]. However, such analyses must be approached with caution, as the lung is not composed of discrete pores, but of interconnected airways structured hierarchically throughout the lung tissue [7]. Thus, while the term “pores” is informative, it remains an artificial representation of the actual lung structure.

Correlative imaging of FFPE tissue is a technique that has gained increasing popularity [8]. Typically, FFPE tissue blocks are first scanned by phase-contrast microCT before sectioning, which allows for spatial registration of all subsequent findings within the 3D context of that modality, as demonstrated by D’Amico et al. [8]. Synchrotron-based phase-contrast microCT is particularly well-suited for this purpose, because it offers enhanced soft-tissue contrast, enabling label-free imaging of standard FFPE blocks, and the high X-ray flux at synchrotron sources permits high-throughput imaging [9]. The “guided cutting” approach uses propagation-based imaging (PBI) to generate detailed 3D images, which guide the sectioning process to specific sites of interest, eliminating the need for labor-intensive serial cutting and thereby improving tissue integrity and reducing workload [10].

In force-spectroscopy–based AFM, the cantilever tip is displaced vertically (along the z-axis) toward and away from the sample at a fixed lateral location, allowing force–distance relationships to be measured without lateral raster scanning, which is characteristic of conventional AFM topographic imaging. The deflection of the cantilever reflects the interaction forces between the tip and the sample surface; these forces are quantified using Hooke’s law, based on the cantilever spring constant and its deflection sensitivity. This method enables a detailed analysis of nanoscale mechanical properties, including stiffness, adhesion, and elasticity. Force-spectroscopy–based AFM has been employed in biomedical applications to assess changes in stiffness (Young’s modulus) across various tissues, including liver, lungs, and brain. It has also been used to study changes in lung tissue stiffness [8].

In summary, our analysis pipeline, integrating phase-contrast microCT, AFM, and histology, revealed significant differences in lung structure and stiffness across the four experimental groups (healthy rats and rats with bleomycin-induced lung fibrosis, both with and without injurious MV). We found that injurious MV led to increased pore volume in healthy control rats, with a more pronounced effect in the absence of pre-existing lung injury. Interestingly, injured regions in bleomycin-treated rats exhibited reduced stiffness following injurious MV, and a spatial correlation between VILI-induced damage and injured regions was observed.

## Results

### Design of the data acquisition and analysis pipeline

To study the relationship between VILI and pulmonary fibrosis in a rat model *ex-vivo*, we designed a comprehensive analysis pipeline, which is outlined in Fig. 1. Lung injury was induced in 8 rats via intratracheal instillation of bleomycin, while 12 healthy rats served as controls. Among the instilled rats, 4 were imaged at baseline by synchortron 4D micro-CT, then underwent injurious MV to induce VILI (bleo-VILI), and were imaged again post-VILI. The remaining 4 animals served as unventilated controls (bleo). The healthy control rats were divided into two groups: 5 were imaged at baseline, then received injurious MV (Con-VILI) and were imaged post-VILI. Seven rats served as unventilated controls (Con). After *in-vivo* strain analysis, the rats were euthanized, and their lungs were harvested. The lungs were then filled with formalin at a constant pressure of 20 cm cmH_2_O and fully fixed in a formalin bath. Subsequently, they were split into multiple regions as described in [4], chemically dried using an ascending ethanol series, and finally embedded in paraffin blocks. Here we used FFPE lung tissue from the upper left lobe (A in Fig. 1) and the lower left lobe (B in Fig. 1). For more detailed information see Fig. 6. These blocks were scanned using phase-contrast microCT to enable both:a) precise targeting of the sectioning process to sites of interest and b) automated 3D characterization of lung structure. Sections of 5 μm thickness were prepared for AFM and histological analysis. The data from all three analysis routes were then integrated using cluster analysis.

**Figure 1.**
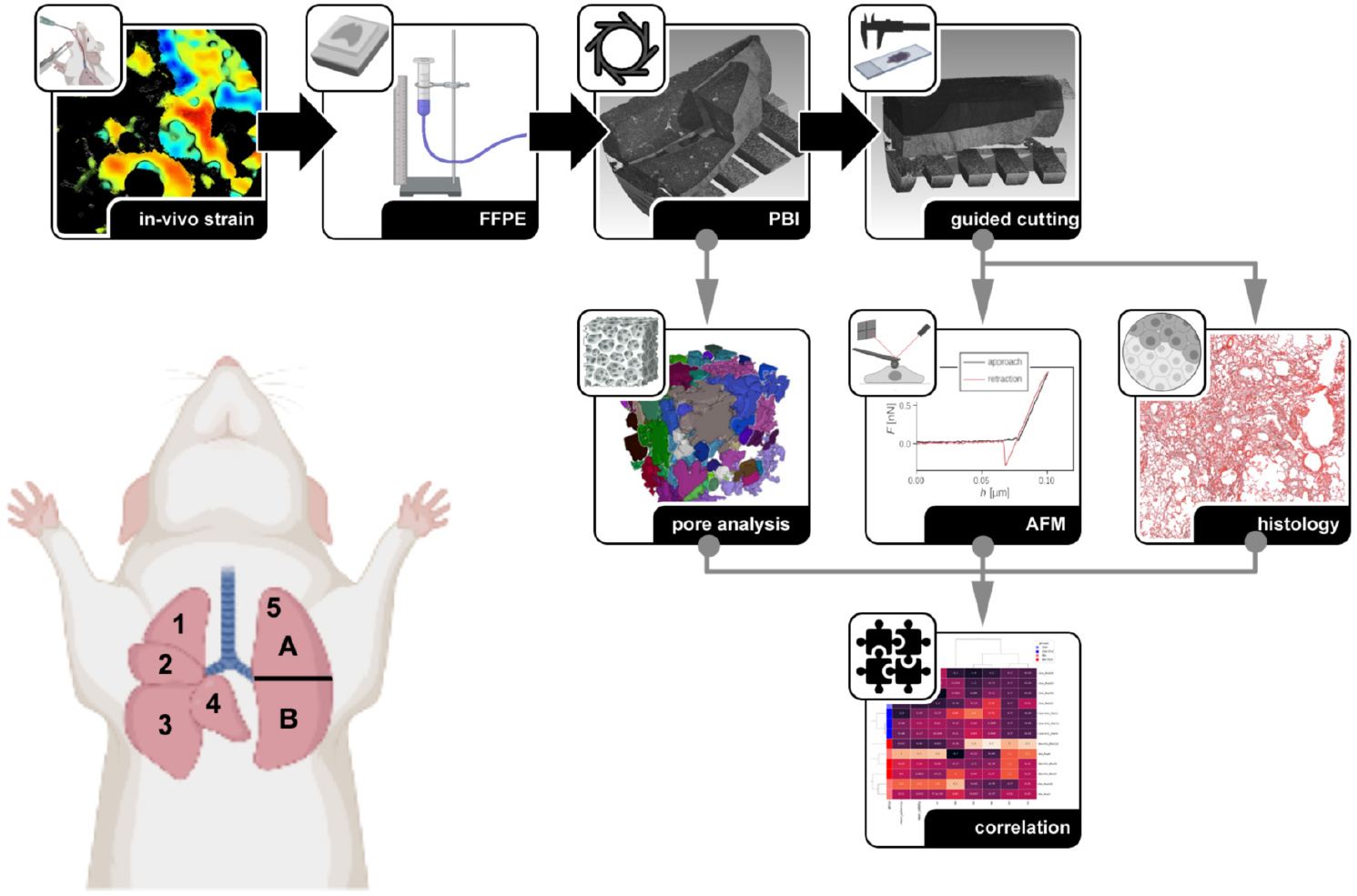
Experimental and data analysis workflow. Following the *in-vivo* experiments, the rats were euthanized, and their lungs were harvested and inflated with formalin at a constant pressure of 20 cm H_2_ O. The rat lung consists of 4 right lobes (superior=1, middle=2, inferior=3, and post caval lobe=4) and one left lobe (left lobe=5). The left lobe was split into two pieces (A=upper part, B=lower part), chemically dehydrated, and embedded in separate paraffin blocks. These FFPE blocks were scanned using PBI, and the resulting data were used to guide sectioning to sites of interest. Pore analysis was conducted on the 3D PBI data. The 5 μm thin tissue sections were subsequently processed for AFM and classical histology. Finally, all the data was used for a comprehensive analysis based on hierarchical clustering.

### Summary of previously reported *in-vivo* measurements

Previously published work from the same cohort demonstrated that bleomycin injury combined with injurious MV induces pronounced alterations in lung mechanics, acinar strain distribution, and collagen nanoarchitecture *in-vivo*. Details can be found in Deyhle et al. [3]. Briefly, at the organ scale, forced oscillation measurements revealed a significant increase in dynamic tissue elastance (H) following bleomycin injury, consistent with elevated global lung stiffness. Injurious mechanical ventilation further modified the mechanical phenotype, but did not restore strain levels to those observed in healthy lungs. Tissue damping (G) showed comparatively smaller changes, and airway resistance was not the dominant contributor to altered mechanics, indicating that parenchymal remodeling primarily governed the observed functional impairment. Four-dimensional synchrotron phase-contrast microCT strain mapping demonstrated that bleomycin-treated lungs exhibited significantly reduced maximal local strain (ε_max_) within aerated acini during tidal ventilation. Importantly, regions of high strain in healthy lungs were spatially redistributed and attenuated in bleomycin-injured lungs, indicating that increased tissue stiffness limited local deformation. Following injurious ventilation, strain heterogeneity increased in control lungs but remained comparatively constrained in bleomycin-treated lungs, suggesting that pre-existing stiffening mitigated excessive regional overdistension. At the nanoscale, collagen fibrillar organization was significantly altered by both bleomycin injury and applied strain. Bleomycin exposure led to increased collagen packing density and structural reorganization consistent with early fibrotic remodeling. Under mechanical strain, collagen fibrils in healthy lungs exhibited measurable nanoscale deformation and reorientation. In contrast, bleomycin-injured lungs demonstrated reduced fibrillar compliance, indicating a mechanically reinforced extracellular matrix. These nanoscale changes correlated with the macroscopic increase in tissue elastance and the reduction in acinar strain.

Importantly, the study demonstrated a direct multiscale linkage: nanoscale collagen remodeling was associated with altered local strain transmission and increased global tissue elastance. Thus, extracellular matrix reorganization after bleomycin injury modifies how mechanical forces are distributed across the lung parenchyma, limiting local deformation during injurious MV. Together, these findings define the *in-vivo* mechanical phenotype of bleomycin-injured lungs as a state of increased global stiffness, reduced acinar strain, and reinforced collagen nanoarchitecture. These results provide the mechanistic basis for interpreting the present *ex-vivo* structural and mechanical measurements and support the concept that extracellular matrix remodeling modulates susceptibility to ventilator-induced lung injury across length scales.

### Results of the pore analysis and force-spectroscopy–based AFM

FFPE lamellae the rat lungs were scanned with PBI at 2 μm resolution. The phase-retrieved and 3D reconstructed volume data sets were analyzed using cubic regions of interest (ROIs) of 300 × 300 × 300 voxel automatically placed with an 100 voxel overlap. ROIs partially including the edge of the reconstructed FOV or ROIs that contained visual air artifacts were discarded. A simple threshold-based segmentation was used to discriminate airways/paraffin from lung tissue. Automatic pore analysis was performed in the airway segment. Since all airways are interconnected, a 3D distance transformation followed by thresholding was applied to separate them into individual *pores*. All pores were analyzed, and their volume, extent (the ratio of the pore to the bounding box), solidity (the ratio of the pore to its convex hull), surface area, and sphericity (the ratio of the surface area of an equal-volume sphere to the actual surface area of the pore) were quantified. This resulted in the analysis of in average 180 pores per region of interest (ROI) and about 685 ROIs per specimens (Con N=7, Con-VILI N=5, Bleo N=4, Bleo-VILI N=4).

Five-micron thick slices were cut from the FFPE tissue of all specimens, with the PBI data guiding the cutting process to target fibrotic regions in the Bleo and Bleo-VILI groups. These slices were then deparaffinized, rehydrated, and analyzed using AFM to probe the alveolar walls (parenchyma) and, where present, the consolidated regions. On average, 30 measurements were taken per region, with 8 regions per slice for each specimen (Con: N=4, Con-VILI: N=3, Bleo: N=3, and Bleo-VILI: N=4). Each individual measurement involved pressing the AFM cantilever with an increasing known force against the tissue, while the bending of the cantilever was monitored by a laser spot. The stiffness, or Young’s modulus, of the underlying tissue was then calculated using the Hertz model [11].

All measurements were averaged for each individual rat and are presented in Fig. 2. Panel a) shows a significantly higher average pore volume in the Bleo group compared to the Con group, as well as a significantly higher pore volume in the Con-VILI group compared to the Con group (Fig. 2a). Panels b) and c) show the results of the histological scoring performed on the specimens. Both the extent of consolidation (consolidation score) and the severity of lung injury (lung injury score) were rated on a 4-point scale from 0 (no presence) to 3 (large area / high severity). Scoring was done on 1024 x 1024 tiles of hematoxylin and eosin (H&E)-stained histological slides from all specimens, with readers blinded to health versus disease status and to preventive versus injurious MV conditions of the specimen. The scores presented are medians for each specimen. Notably, no consolidation was observed in either of the control groups (Con, Con-VILI). However, lung consolidation was more severe in the Bleo-VILI group than in the Bleo group, with one Bleo specimen receiving a consolidation score of 0. (Fig. 2b). Panel c) highlights the difficulty in distinguishing the effects of VILI from the lung damage caused by progressive inflammation and fibrosis. Comparing both Bleo and Bleo-VILI groups, the Bleo-VILI group showed a trend toward higher lung injury scores. In contrast, both both Con and Con-VILI groups showed virtually no signs of lung injury (Fig. 2c), which contradicts the results of the pore analysis shown in Fig. 2a). Fig. 2d) demonstrates that no significant differences in the stiffness of the alveolar walls in the lung parenchyma were found between all 4 groups. In the consolidated regions (Fig. 2e) a trend of reduced stiffness in presence of VILI was observed in the Bleo-VILI group compared to the Bleo group.

**Figure 2.**
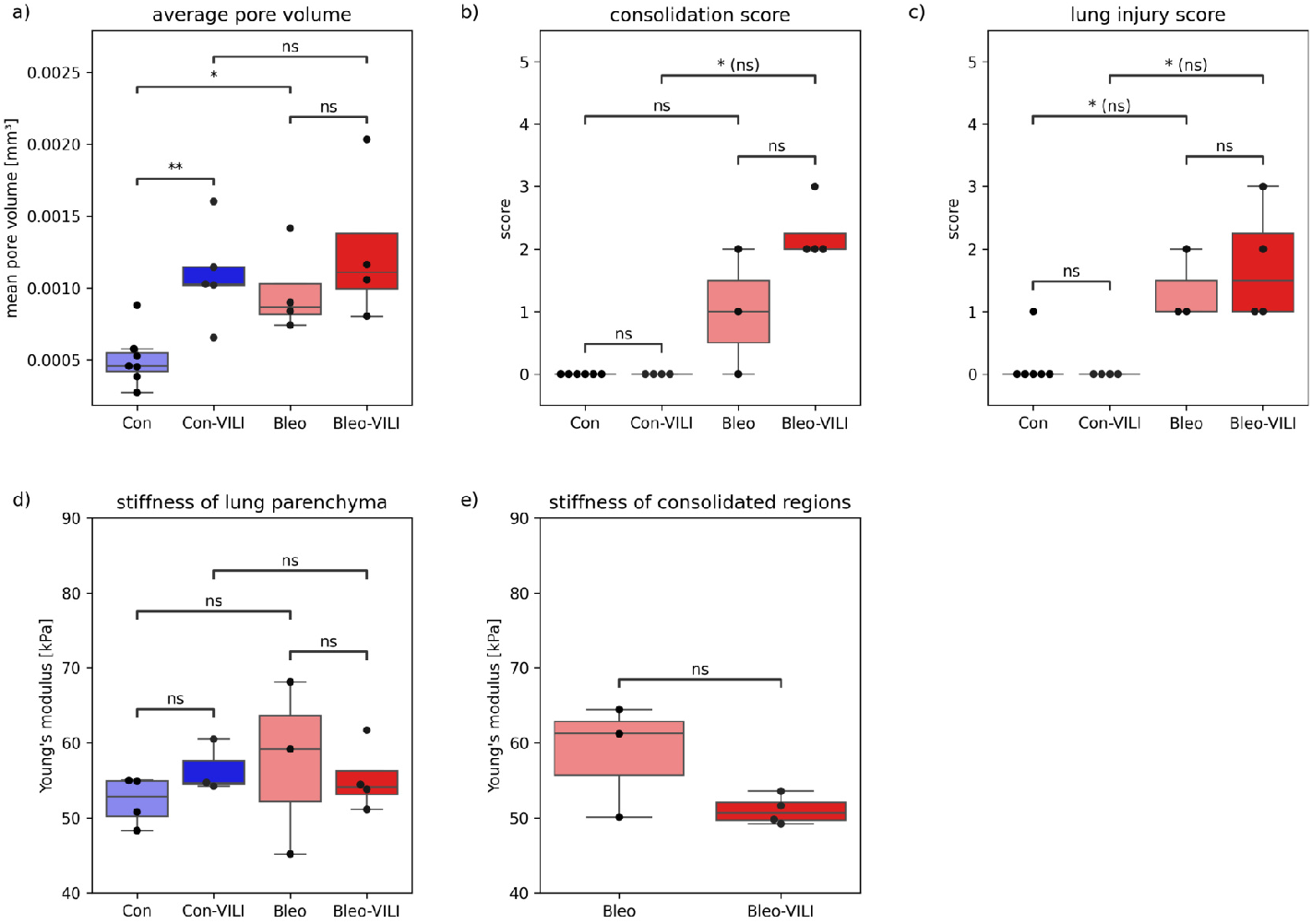
Quantitative results. a) A significant increase in average pore volume was observed between healthy controls without VILI (Con) and those with VILI (Con-VILI), as well as between controls (Con) and bleomycin-induced lung fibrosis (Bleo). b) and c) Histological scoring revealed increased consolidation in the Bleo group, with even higher consolidation observed in the Bleo-VILI group, as well as greater lung injury in both Bleo and Bleo-VILI groups, compared to their respective control groups. Interestingly, no increase in the lung injury score was found in the Con-VILI group compared to the Con group, highlighting the challenges of assessing this aspect using two-dimensional (2D) histology. d) AFM measurements of alveolar regions indicated a stiffening of the lung parenchyma in the Con-VILI group compared to the Con group. e) In contrast, a reverse trend was observed in the stiffness of consolidated regions in the Bleo and Bleo-VILI groups.

The relation between *in-vivo* global respiratory elastance (H) [3] and mean pore volume is shown in Fig. 3. Both parameters changed in the same direction in response to injurious mechanical ventilation, both in initially healthy lungs and in early bleomycin-induced lung injury. Bleomycin injury also independently increased both respiratory tissue elastance and mean pore volume.

**Figure 3.**
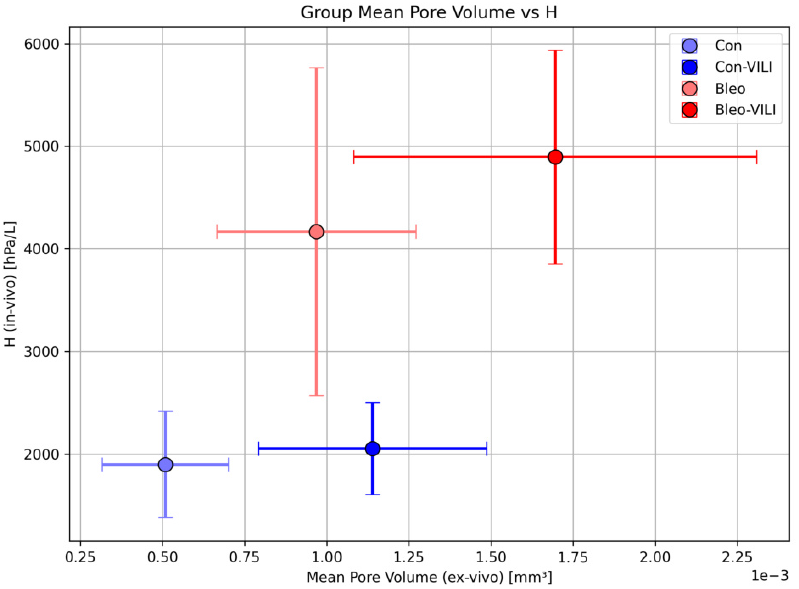
Relation between global lung elastance and pore volume. Global respiratory elastance (H) [3] and pore volume both increased in healthy controls (Con) as a result of injurious mechanical ventilation (Con-VILI). Both parameters were significantly elevated in bleomycin-induced lung fibrosis (Bleo) vs healthy controls, and increased further in response to injurious mechanica vnetilation (Bleo-VILI).

To assess whether the specimens from the 4 different groups exhibit unique combinations of features derived from pore analysis and AFM, hierarchical clustering was performed using z-scoring, cosine-similarity, and averaging as linking criteria (Fig. 4). The results clearly distinguish non-fibrotic from fibrotic specimens. The impact of VILI on lung damage, manifested as larger pore sizes, was more consistently observed in the non-fibrotic groups, which aligns closely with the pore analysis findings. However, the clustering was primarily influenced by histological scoring, as evidenced by the histological examples in Fig. 4. These images highlight striking differences in the presence of consolidation and the extent of damage across the 4 groups. Interestingly, the presence of VILI in the not bleomycin treated control group (CN-VILI) was not picked up during the scoring process, highlighting the advantage of 3D pore analysis.

**Figure 4.**
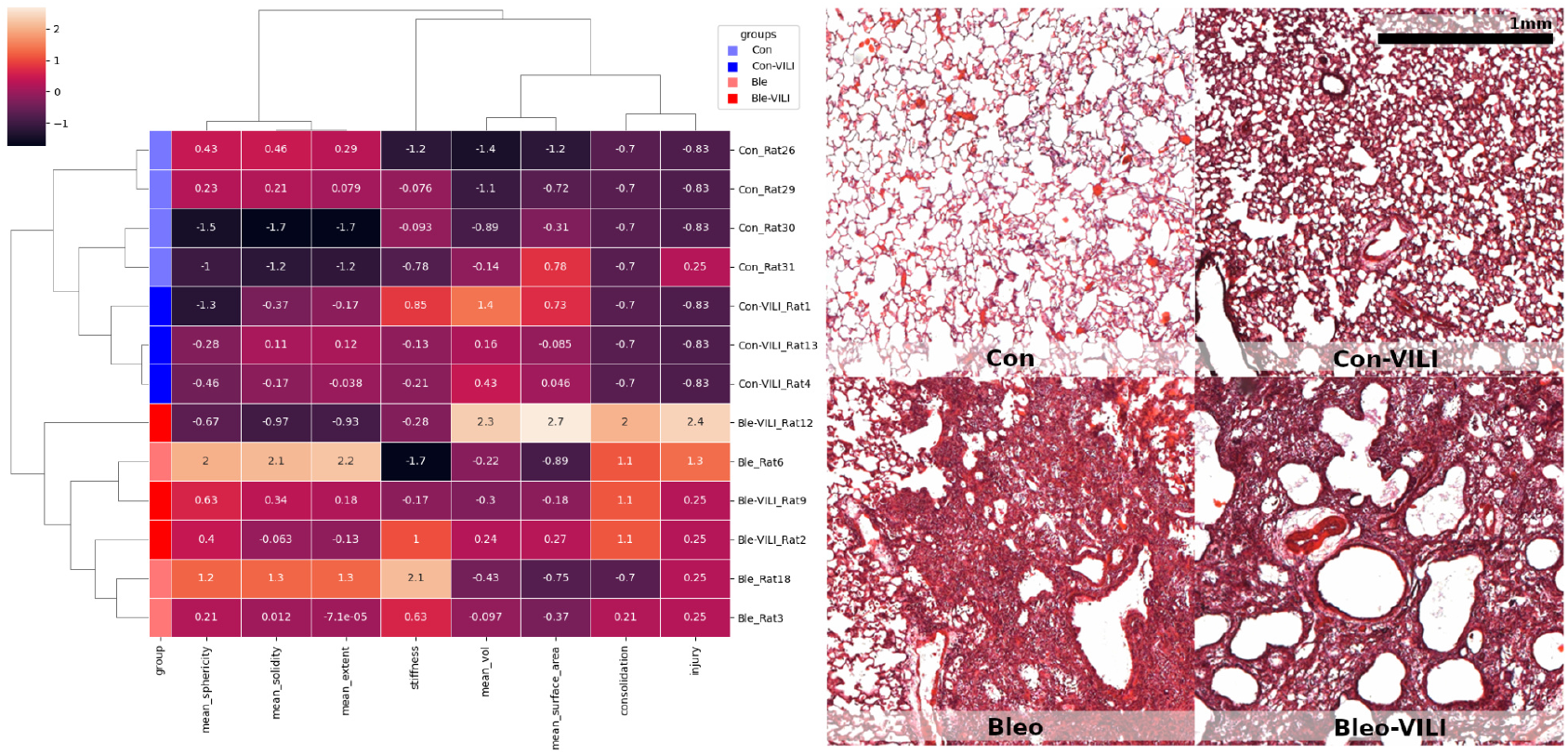
Cluster analysis. The clustermap clearly distinguishes between healthy controls with VILI inducing mechanical ventilation (Con-VILI), healthy controls with protective mechanical ventilation (Con), and fibrotic specimens (Bleo, Bleo-VILI). Notably, the separation between VILI inducing MV and protective MV subjects is more pronounced in the control groups (Con vs. Con-VILI), which aligns with the results of the pore analysis. However, the clustering is primarily driven by the histological features of the ‘consolidation’ and ‘injury’ scores. This is further supported by the histological analysis, which reveals distinct differences in the amount of consolidation and extent of tissue damage across the four groups.

### Correlation and spatial integration of the results

A key feature of our analysis pipeline is the ability to co-register the results of subsequent two-dimensional (2D) measurements—AFM and histology—with the 3D PBI datasets of uncut FFPE tissue. To address the deformations introduced during the sectioning process, we employed elastic registration techniques as described by D’Amico et al. [8]. In the first step the virtual cutting plane in the PBI data set was identified that corresponded to the plane in which the histological sections were cut. Deformation was confined to the cut sections rather than the entire FFPE tissue block. This allowed us to apply a straightforward 2D-2D elastic registration once the virtual cutting plane was defined. The result of this registration is shown in Fig. 5a) as a checker-board visualization, where a high-quality alignment is evident from the continuous tissue structures across both datasets. Importantly, both the virtual PBI plane and the histological image were cropped to the same rectangular region to ensure a precise overlay. This alignment enabled the successful comparison of 3D structural alterations from the PBI data with corresponding histological features (Fig. 5b), providing valuable insights into tissue architecture.

**Figure 5.**
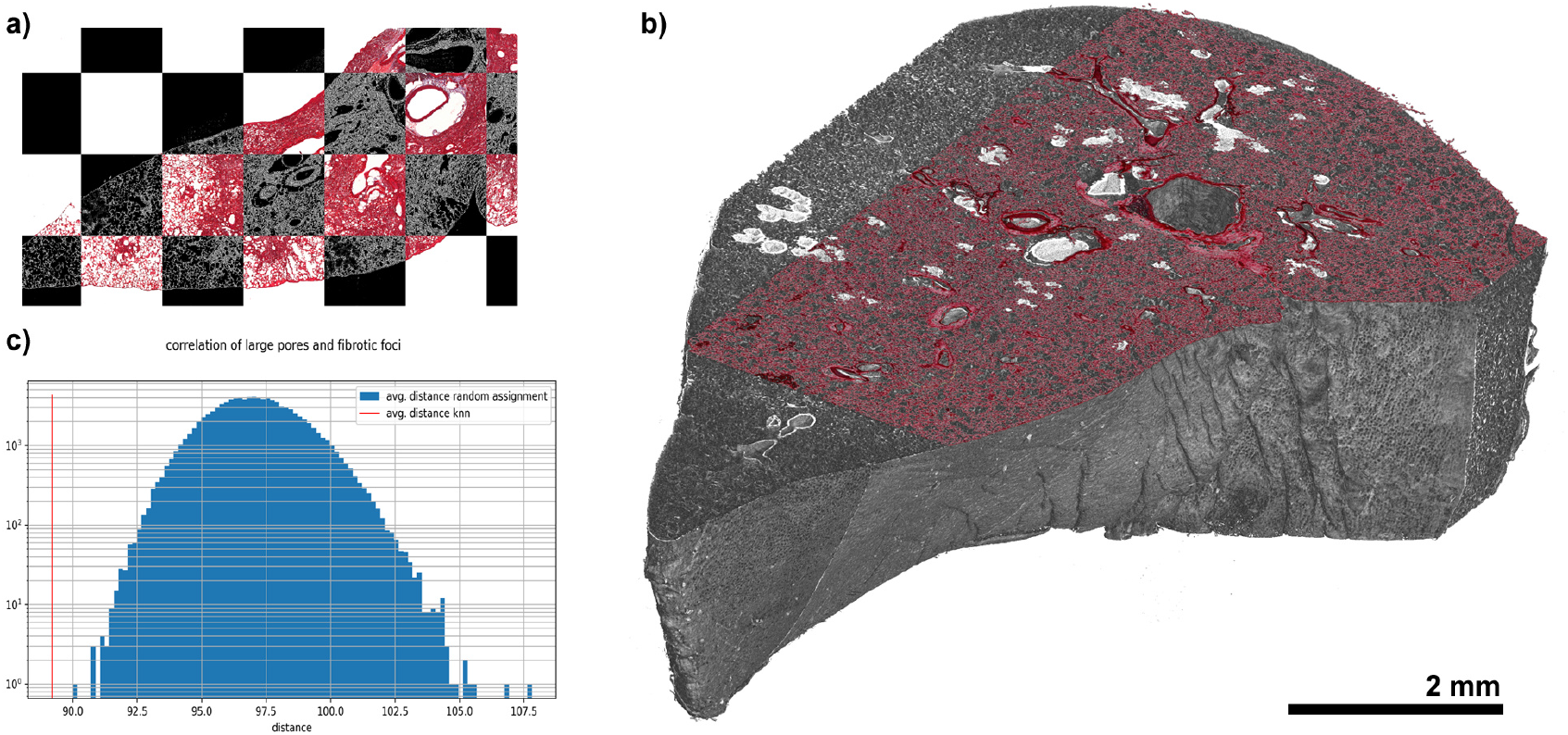
Spatial integration and correlation of Bleo-VILI lungs. a) Checkerboard view of the elastic registration between a plane isolated from a PBI dataset (black background) and a subsequently generated H&E-stained section of the same region, demonstrating a high precision of the match. b) 3D rendering of the PBI dataset of a control lung, integrated with the registered histology (note: the histology has been cropped for improved visibility). c) Result of the k-nearest neighbor (kNN) analysis of fibrotic foci and VILI regions. The average distance (indicated by the red line) is significantly smaller than in all bootstrap tests, suggesting a spatial relationship between fibrotic regions and VILI.

**Figure 6.**
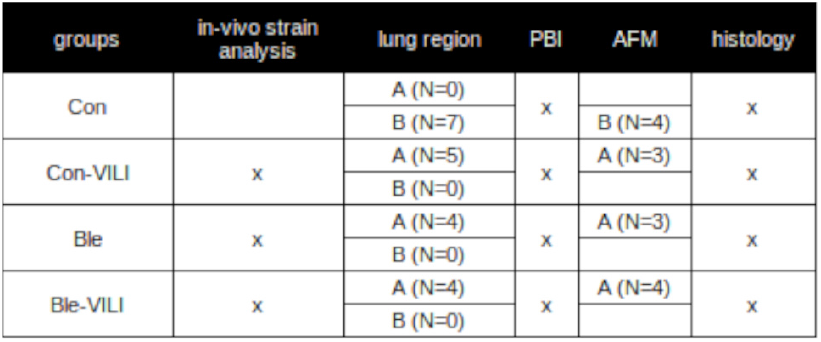
Overview of the study design. For *ex-vivo* analyses, the left lobe of the rat lung was divided into two pieces (A=upper part, B=lower part). Con=Healthy control rats after protective mechanical ventilation. Con-VILI=Healthy control rats after injurious mechanical ventilation. Bleo=Bleomycin-induced lung injury rat model after protective mechanical ventilation. Bleo-VILI=Bleomycin-induced lung injury rat model after injurious mechanical ventilation. PBI=Propagation-based imaging. AFM=Atomic force microscopy.

The primary goal of the study was to investigate whether there is a spatial relationship between lung regions damaged by VILI and fibrotic areas. To this end, a distance transformation of the lung tissue was performed, followed by a thresholding at 10 voxels, which resulted in isolated regions of thickened lung tissue. These regions were then represented by their centroids using a blob-analysis algorithm. The positions of these centroids constituted the first point cloud (1).

From the pore analysis described above, the centroids of pores with a volume greater than 0.00064 mm^3^ were extracted to form the second point cloud (2).

A 10-nearest neighbor analysis was performed for each point in set 1 with respect to set 2, and the average distance between points was recorded. To assess the statistical significance of this distance, we implemented a bootstrap procedure, repeating the process 100,000 times. In each iteration, new point clouds, denoted as 1′ and 2′, were randomly generated by resampling with replacement, while maintaining the same cardinality as the original sets. The 10-nearest neighbor analysis was then repeated for these resampled point clouds, and the results were compared to those of the original point clouds. The p-value was calculated as follows:

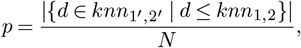

where *knn*_*i,j*_ represents the average distance in the *k*-neighborhood of each point in set *i* with respect to set *j*, and *N* is the total number of bootstrap iterations.

We found a significantly lower average distance for larger pores in the 10-nearest neighbor analysis of fibrotic foci compared to the bootstrap analysis (Fig. 5c). This suggests a functional correlation between regions of fibrosis and VILI.

## Discussion

In this study, we established a comprehensive multiscale pipeline for the quantitative analysis of lung porosity and mechanical stiffness in FFPE rat lungs. By combining high-resolution, label-free phase-contrast microCT, atomic force microscopy (AFM), and classical histology, we investigated the interplay between ventilator-induced lung injury (VILI) and bleomycin-induced inflammation and early fibrosis. Automated pore analysis, image registration, and cluster analysis enabled objective quantification of structural and mechanical features across spatial scales.

Our results demonstrate a significant increase in average pore volume, indicative of microscale airspace expansion, both following injurious ventilation in initially healthy lungs and after bleomycin-induced inflammatory injury with early matrix remodeling. Notably, airspace enlargement was more pronounced in healthy lungs exposed to injurious ventilation than in fibrotic lungs.

Post-mortem histology from the pre-protective ventilation era revealed frequent bronchiolar dilatation and alveolar overdistension in ARDS patients ventilated with high tidal volumes and pressures, resembling emphysematous changes [12]. Similar findings have been reproduced experimentally in mechanically ventilated pigs with induced pneumonia [13, 14]. Airspace enlargement has also been described in models of chronic inflammation, including endotoxemia [15], oxygen toxicity [16], and starvation [17]. However, whether mechanical stress alone can induce persistent airspace enlargement has remained controversial [12].

Our data show that early bleomycin-induced lung injury (7 days) was likewise associated with airspace enlargement. This may reflect inflammatory elastin degradation, as neutrophil elastase released during type I inflammation can degrade alveolar elastin and promote septal destruction [18, 19]. Alternatively, matrix deposition and stiffening during early fibrosis may increase radial traction forces on alveolar walls, promoting mechanical dilation in a manner analogous to retraction bronchiectasis in fibrotic lung disease [20]. These mechanisms are not mutually exclusive and may coexist during early remodeling.

Importantly, our findings further suggest that high-strain mechanical ventilation can induce microscale airspace enlargement even in initially healthy lungs. While extreme ventilatory pressures can produce overt structural failure such as interstitial emphysema and pneumothorax [21], more subtle matrix-mediated mechanisms are likely operative at subclinical strain levels. The extracellular matrix transmits mechanical forces to resident cells, activating mechanotransductive pathways that may amplify inflammatory responses [22]. Qualitative histology revealed inflammatory cell infiltration and matrix thickening in ventilated lungs, consistent with this interpretation. Together with the concomitant increase in global respiratory elastance, these findings suggest that altered matrix mechanics and inflammation, rather than acute septal rupture, underlie the observed microscale enlargement. Notably, these structural changes are not detectable clinically or by conventional CT imaging.

Our results align with recent nanoscale studies demonstrating strain- and bleomycin-dependent remodeling of collagen architecture in the same animals [3]. By linking global *in-vivo* mechanical alterations to microscale airspace enlargement and nanoscale matrix remodeling, our data provide a coherent multiscale framework connecting organ-level mechanics to extracellular matrix structure (Fig. 2).

Phase-contrast microCT, combined with guided sectioning, allowed precise localization of inflamed and fibrotic regions, and enabled quantitative pore analysis in FFPE tissue [8, 23]. The use of single distance phase retrieval [24] enhanced soft-tissue contrast, while AFM force spectroscopy provided local stiffness measurements at the alveolar wall level [8]. Importantly, our correlative approach integrates 3D structural data with 2D AFM and histology, mitigating the limitations of con-ventional histology, which may underestimate VILI in healthy lungs due to subtle or sectioning-induced artifacts [23], and in heterogeneous lungs due to sampling bias [25].

The use of automated pore analysis offers a sensitive metric for detecting structural damage induced by MV. Pore metrics, such as volume, extent, and sphericity, provide complementary insights to histological scoring, capturing changes that may not be visually evident. Such an approach has for instance been used by Scott et al. [26]. In our study, pore analysis detected significant VILI in healthy lungs that was not reflected in histology, demonstrating the advantages of high-resolution, 3D quantification. Nevertheless, it is important to recognize that “pores” in the lung are an abstraction of the hierarchical airway network [7], and segmentation of FFPE tissue introduces potential biases, such as artificial cuts or undersampling.

Several limitations warrant consideration. First, FFPE tissue alters mechanical properties due to formalin-induced cross-linking [27], and AFM measurements on FFPE are generally lower in sensitivity than on fresh tissue [28]. Second, rat lungs were too large to embed intact in standard paraffin blocks, requiring dissection into upper and lower lobes. Consequently, PBI scans captured only parts of the lung, preventing normalization of pore metrics to total lung volume. Third, air inclusions in FFPE specimens produced artifacts in PBI scans, necessitating selective ROI analysis [29]. Only the Bleo-VILI specimens originated from the identical animals subjected to *ex-vivo* analysis in the present study. Consequently, direct *in-vivo*–*ex-vivo* correlations could be performed only for this cohort. Finally, the bleomycin model exhibits spatial heterogeneity in fibrosis, which complicates comparisons between groups [30].

Despite these limitations, our results are consistent with the *in-vivo* observations of reduced VILI in fibrotic lungs [31]. The integration of nanoscale observations from Deyhle et al. [3] further supports the notion that collagen remodeling during fibrosis mitigates additional alveolar strain. Moreover, our k-nearest neighbor analysis revealed a spatial correlation between fibrotic regions and large pore formation, suggesting that pre-existing structural heterogeneity modulates the pattern of injury induced by MV.

Overall, this study highlights the advantages of an automated, correlative 3D analysis pipeline for assessing lung injury and fibrosis, while emphasizing the importance of standardized sample preparation and careful handling of FFPE artifacts. The combination of phase-contrast microCT, AFM, and histology provides a multi-scale perspective on lung structure and mechanics, enabling the detection of subtle VILI effects that may be overlooked with conventional 2D histology.

## Conclusion

We implemented a multiscale correlative pipeline integrating 3D label-free phase-contrast microCT, automated pore analysis, atomic force microscopy, and classical histology to characterize structural and mechanical alterations in lung tissue.

Our data demonstrate that both injurious mechanical ventilation and bleomycin-induced inflammatory injury independently promote microscale airspace expansion. The comparison with global respiratory mechanical changes suggests that the observed airspace expansions are induced by parenchymal stiffening and enhanced retraction forces rather than direct mechanical rupture. These structural changes bridge a critical spatial and conceptual gap by linking global *in-vivo* mechanical alterations to nanoscale collagen remodelling previously demonstrated in the same animals [3]. In doing so, this work connects organ-level mechanical dysfunction to extracellular matrix remodeling across length scales.

Spatial analysis further revealed co-localization of fibrotic regions and ventilation-induced structural damage, supporting the concept that pre-existing matrix heterogeneity modulates stress distribution and injury propagation. Together, these findings provide a mechanistic multiscale framework for understanding the interaction between fibrosis and ventilator-induced lung injury.

## Conclusion

In this study, we successfully implemented a comprehensive multiscale analysis pipeline for lung tissue, integrating 3D label-free phase-contrast microCT, automated pore analysis, atomic force microscopy, and classical histology. Our data demonstrate that increased mechanical stress resulting from injurious mechanical ventilation, as well as inflammatory lung injury induced by intratracheal bleomycin instillation, independently promote airspace expansion at the microscale. These microscopic structural changes link the global \textit{in-vivo} alterations in lung mechanics to nanoscale structural remodelling of collagen previously observed in the same animals [3]. In doing so, our study connects mechanical dysfunction at the organ level with matrix-level structural changes across spatial scales. Furthermore, spatial analysis revealed a significant co-localization of fibrotic regions and VILI-induced airspace damage, emphasizing the role of mechanical stress concentration in injury propagation. This multiscale perspective provides a mechanistic framework for understanding the interplay between fibrosis and VILI. By correlating *ex-vivo* microstructural and mechanical parameters with *in-vivo* lung function, we show that nanoscale tissue stiffness, pore geometry, and consolidation patterns are closely associated with organ-level mechanics, including elastance, tidal volume, and airway resistance.

## Methods

### Experimental setup, sample preparation

The experiments were performed on 20 Sprague-Dawley rats, average weight: 399±26 g. The animals were divided into 4 groups (see Fig. 6): Controls, with protective MV (Con); Controls after ventilator-induced lung injury induction (Con-VILI), Bleomycin-induced lung injury with protective MV (Bleo), and bleomycin after induction of VILI (Bleo-VILI). In the latter groups, the animals received a single intratracheal dose of bleomycin (Sigma-Aldrich, St. Louis, MO, United States), with a concentration of 1000 iU in 200 μl saline. Control animals received the same volume of saline only. 14 animals underwent *in-vivo* synchrotron X-ray microscopy imaging for a separate study. After baseline image acquisition, injurious MV was initiated by increasing peak respiratory pressure to 41±2 cm H_2_O, 0 PEEP, and respiratory rate was reduced to 30 bpm for 20 minutes to induce VILI. At the end of the *in-vivo* imaging, the animals were euthanized by intraperitoneal injection of pentobarbital sodium (Dolethal, 200 mg/kg, Vetoquinol, Lure, France) and the heart and lungs were dissected and removed en bloc for histological analysis. The left lungs were fixed in 4% paraformaldehyde at 20 cm H_2_O, dehydrated with a graded ethanol series, split in an upper part (A in Fig. 1) and a lower part (B in Fig. 1), and embedded in paraffin (Fig. 6).

### Phase-contrast microCT

All FFPE tissue specimens were scanned at the SYnchrotron Radiation for MEdical Physics (SYRMEP) beamline [32] using a white/pink beam in propagation-based phase-contrast mode with a 150 mm sample-to-detector distance. The beam was filtered through a 0.5 mm silica foil, resulting in a mean energy of 16.7 keV. 360° off-center scans were performed with 3600 projections and a 50 ms exposure time per projection, using a water-cooled Orca Flash 4.0 sCMOS detector (2048×2048 pixels) coupled to a 17 μm Gallium Gadolinium Garnet (GGG) scintillator. The pixel size was 2 μm, giving a reconstructed field of view of approximately 7×4 mm^2^. To capture the entire FFPE lung specimen, 2–4 scans were performed with a vertical offset of 3.5 mm. The slices were reconstructed using SYRMEP Tomo Project (STP) software [33], with the TIE-Hom phase retrieval algorithm [24] (d/b ratio of 100) applied before filtered back-projection. Vertical slices for each lung were then stitched together using a custom-made Python script to create a complete 3D representation of the specimen.

### 3D pore analysis for characterizing anatomical alterations

PoreSpy [34] was used for quantification of the airway network. To this end a binarization of 300 x 300 x 300 ROIs within the PBI data set was performed using a threshold value of 55, followed by morphological binary fill-holes and closing operations. To split the connected airways into individual pores, a distance transform was applied on the segmented airways combined with a threshold of 10, followed by a label-watershed segmentation. For the final segmented isolated pores the following metrics were calculated:

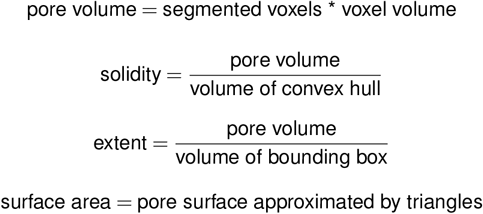

### Guided sectioning, histological staining, sample preparation for AFM

The 3D PBI datasets were analyzed to investigate structural changes and identify fibrotic regions, a critical step for precisely targeting the fibrotic areas during paraffin block sectioning. The 3D lung volumes were converted to 8-bit format and loaded into VGSTUDIO MAX (Volume Graphics, Germany) for 3D visualization and identification of pathological regions. Initially, the data were aligned to the surface of the paraffin block. After calculating the depth of the ROIs based on the PBI microCT data, each paraffin block was sectioned at this depth using a microtome. Additionally, the cutting plane within the 3D rendering of the PBI data was stored for future reference. Given that cutting was performed on cooled FFPE tissue while the PBI scans were conducted at room temperature, a 20% shrinkage correction factor was empirically determined and applied to adjust the cutting positions. To facilitate spatial registration of both AFM and histological data to the PBI dataset, 3 to 5 adjacent 5-μm thick sections were prepared. The first section was stained with H&E for reference, while the second was mounted on a glass slide without a cover slip to enable AFM measurements in parallel with imaging using an inverted microscope integrated into the AFM system.

### AFM measurements

To prepare for biomechanical characterization, the slide was deparaffinized and rehydrated using the following protocol: 1 hour at 65°C, 15 minutes in xylol, rinse in fresh xylol, 10 minutes in 99% ethanol, 10 minutes in 96% ethanol, 5 minutes in 50% ethanol, and finally transferred to water.

AFM measurements were performed using a Nanowizard 4 (JPK, Bruker Nano GmbH) on an Olympus IX83 inverted microscope. The measurements were conducted in force spectroscopy mode with a 10 nN set point and a Z speed of 2 μm/s, using a pre-calibrated SiO_2_ cantilever with a spring constant of 0.062 N/m and a 2.5 μm bead. All experiments were conducted in water. Young’s moduli were retrieved using JPK Data Pre-processing software (version 7.0.165), with baseline subtraction and contact point determination applied to the force curves before elasticity fitting. To model the tip-sample interaction and extract biomechanical data the Hertz model was used [11]. This model assumes the sample is elastic and homogeneous, and the use of a non-deformable spherical tip. A custom-made Python script was used to overlay the force curves on the corresponding microscopic images and to convert them into 3D bar plots to enable spatial registration with the PBI data as introduced by D’Amico et al. [8].

### Histological analysis

All H&E-stained sections were imaged at 5x magnification using a mosaic scan in an Axioskop microscope (Zeiss) equipped with a digital camera (Micropublisher 5.0, QImaging Surrey) to capture the entire crosssection of the specimen. The resulting images were divided into 1024 x 1204 px^2^ tiles. Tiles with less than 25% tissue content were excluded from analysis. The remaining tiles were then randomized and assessed for the extent of consolidation and lung injury, each scored using a 4-point scale ranging from 0 (not present) to 3 (strong effect). Readers were blinded to health versus disease status and to preventive versus injurious MV conditions of the specimen. The median score for each specimen was used for subsequent statistical analysis.

### *In-vivo* respiratory mechanics and strain mapping

The *in-vivo* datasets summarized and correlated in the present study were previously published in detail by Deyhle Jr. et al. [3] and are included here to enable correlation with the newly acquired ex-vivo structural and mechanical measurements.

For the *in-vivo* component, respiratory mechanics and local lung strain data were obtained from mechanically ventilated Sprague–Dawley rats as described previously [3]. In brief, male Sprague–Dawley rats (average weight: 399 ± 26 g) were anesthetized, tracheostomized, and mechanically ventilated under controlled conditions. Animals were studied at baseline and after a short injurious MV protocol designed to induce VILI. Injurious MV was applied using high tidal volumes (≈ 65 ml/kg) with Pip of ≈ 41 cm H_2_O at zero PEEP, consistent with established experimental VILI models [3].

Respiratory mechanics were assessed using the forced oscillation technique (FOT) during MV, yielding parameters of global lung function including tissue elastance (H), tissue damping (G), and airway resistance (Raw). Oscillatory measurements were acquired over a range of frequencies and analyzed to characterize the viscoelastic behavior of the respiratory system under both baseline and injurious MV conditions [3]. Dynamic tissue elastance (H) was used as the primary metric of global lung stiffness in the correlation analyses presented in the current study.

To obtain spatially resolved measures of lung deformation, 4D *in-vivo* phase-contrast microCT was performed at a synchrotron X-ray source, using retrospective gating on the continuously acquired projection images. High-intensity X-rays were rendered quasi-monochromatic and recorded using a high-resolution detector, yielding an isotropic voxel size of approximately 6 μm. This enabled dynamic imaging of deep pulmonary acini during tidal ventilation [3]. Using a stepwise non-rigid image registration algorithm, quantitative maps of local microscopic lung tissue strain (ϵ) were computed within aerated regions throughout the respiratory cycle, allowing determination of maximal local strain (ϵ_max_) for each condition in various volumes of interest throughout the FOV of the entire lung.

These *in-vivo* datasets of respiratory mechanics and local strain form the basis for correlation with the *ex-vivo* structural and mechanical parameters derived from phase-contrast microCT pore analysis and AFM in FFPE lung tissue. The combination of dynamic strain imaging and global respiratory mechanics provides complementary perspectives on lung function, with local deformation metrics reflecting tissue-scale mechanical behavior and FOT parameters capturing organ-scale viscoelastic properties [3].

Importantly, only the Bleo-VILI specimens correspond to the identical animals analyzed *ex-vivo* in the present study. Consequently, direct *in-vivo*–*ex-vivo* correlations were restricted to this cohort.

### Summary of the study design

To summarize the complex study design all relevant information are shown in Figure 6.

### Software and statistics

Statistical analysis was performed with Seaborn (0.13.2) [35] in combination with statannotations (0.7.1) [36]. For comparison of difference in the median between groups the Mann-Whitney-Wilcoxon test two-sided with Benjamini-Hochberg correction for multiple testing was used. A p < 0.05 was considered statistical significant. P-values are indicated on the figures as (not significant (ns): 0.05 < p <= 1.00, *: 0.01 < p <= 0.05,**: 0.001 < p <= 0.01, ***: 0.0001 < p <= 0.001,****: p <= 0.00001, * (ns) indicates significance for raw p-values and not significant after Benjamini-Hochberg correction).

To analyze a spatial correlation between fibrotic regions and injured lung regions caused by VILI, a kNN analysis between two 3D point clouds (centroids of fibrotic regions and centroids of large pores) was performed using scikit-learn aka sklearn (1.6.0) [37].

For the comparison of *ex-vivo* and *in-vivo* results the Spearman correlation in scipy (1.16.3) [38] was used. Figures have been generated with seaborn, matplotlib (3.10.0), and Photoshop 6.0 (Adobe Systems Incorporated (2000)).

## Additional Information

### Ethics

The care of animals and the experimental procedures were in accordance with the Directive 2010/63/EU of the European Parliament on the protection of animals used for scientific purposes and complied with the ARRIVE guidelines. Experimental procedures were evaluated and approved by the local institutional ethical review board and the French Ministry of Higher Education and Research (authorization number: APAFIS#31021-2021040617424365).

## Acknowledgements

The authors thank Sarah Garbode (Clinic of Haematology and Medical Oncology, UMG Goettingen), Bärbel Heidrich and Regine Kruse (Translational Molecular Imaging, MPI-NAT Goettingen) for excellent technical support in sample preparation, sectioning, and staining. We once again thank the entire team of the SYRMEP beamline of the Italian synchrotron for their support of the phase-contrast microCT acquisitions.

## Author contributions

C.D., S.B., S.K., L.O., and F.A. designed the experiment. C.D., M.S., L.D., R.M., and G.T. performed the *ex-vivo* experiments. R.D., L.F., I.P.M., J. C.-P., G.P., and S.B. performed the *in-vivo* experiment. M.S., L.D., C.V.B., C.D., and S.B. analyzed the data. M.S., L.D., C.V.B., C.D., and S.B. wrote the manuscript. All authors have approved the final version of the manuscript for publication.

